# Whole-fingertip 3D Skin Surface Deformation under Tangential Loading

**DOI:** 10.64898/2026.04.15.718652

**Authors:** Donatien Doumont, Philippe Lefèvre, Benoit P Delhaye

## Abstract

During tactile interaction, skin deformation drives the widespread activation of tactile afferents distributed across the fingertip. Yet the full spatial extent and evolution of these deformations remain largely unquantified. Using high-resolution 3D imaging, we reconstructed the complete volar surface of the fingertip under progressive tangential loadings typical of object manipulation. We show that much of the deformation occurs in the out-of-contact regions, accounting for approximately 70% of the total deformation energy. This deformation consistently initiates in the peripheral zones and smoothly propagates inward as partial slip develops. Tangential loading also induces pronounced directional asymmetries and local curvature changes, reflecting both surface and bulk tissue deformation. Furthermore, we observe localized strain patterns consistent with skin wrinkling across all participants, with individual variations in intensity and location driven by distinct frictional and biomechanical properties. This dataset provides a strong foundation for developing highly accurate biomechanical models and for linking fingertip mechanics to tactile neural encoding.

## Introduction

Our fingertips play a central role in object manipulation, serving as a formidable sensory organ that enables stable and precise grasping (Westling & Johansson, 1987; Augurelle *et al*., 2003). During object manipulation, normal and tangential forces applied at the contact interface induce skin deformation, thereby activating a large population of tactile afferents distributed across the fingertip (Devecioğlu *et al*., 2026). These neural responses provide us with rich information about contact stability, friction, and object properties. Although numerous studies have demonstrated that stimulus features are reliably encoded at the population level (Jenmalm *et al*., 2003; Johansson & Birznieks, 2004; Saal *et al*., 2009; Khamis *et al*., 2015; Delhaye *et al*., 2019), individual tactile afferent signals cannot be interpreted without accounting for the deformation of the skin (Johansson & Flanagan, 2009; Delhaye *et al*., 2021*a*).

Because tactile afferent endings are embedded in the skin, afferent responses do not provide a direct copy of a tactile stimulus but instead reflect the deformation field of the skin. Fingertip deformation depends on its geometry and on the complex biomechanical properties of the skin, including hyperelasticity, viscoelasticity, anisotropy, and multilayered structure (Serina *et al*., 1997; Serhat & Kuchenbecker, 2021; Duprez *et al*., 2024). These properties determine how contact forces are distributed and transmitted to mechanoreceptive endings. For instance, due to its viscoelastic nature, prior stimulation can alter the mechanical state of the skin and thereby shape subsequent afferent responses (Saal *et al*., 2025). Furthermore, the large variability in skin stiffness and thickness across individuals further obscures the direct interpretation of afferent activity when pooled together on a generic fingertip (Birznieks *et al*., 2001; Wheat *et al*., 2010). At the moment, the best afferent models rely on much simplified mechanics and geometry, thus failing to explain the responses to stimuli and force levels that imply large and spread-out deformations, such as those implied by object manipulation (Saal *et al*., 2017).

Advances in optical imaging have enabled direct measurements of the fingertip deformation (Li *et al*., 2020; Kao *et al*., 2022; Bertsch *et al*., 2025; Corniani *et al*., 2025). At the surface of the skin, strains, i.e. local deformation, of large amplitudes were observed (up to 50%), and strain patterns were highly dependent on the stimulus direction or orientation (Delhaye *et al*., 2016, 2021*b*; Schiltz *et al*., 2022; de Dunilac *et al*., 2023). Furthermore, strain waves were tightly coupled with the propagation of partial slip and possibly signaling contact stability (Delhaye *et al*., 2024). However, those studies were inherently restricted to measuring skin deformation either within the contact interface or in a limited region, while deformation extends well beyond the contact region.

We recently developed a method to reconstruct the evolution of the 3D surface shape of the fingertip skin under mechanical loading (Doumont *et al*., 2025). Even for the simplest mechanical interactions with the environment, the contact initiation revealed complex strain waves following the moving contact border as loading increased. In the present study, we quantified how the entire surface of the fingertip deforms during a tangential loading that induces progressive slip. We aimed to characterize how deformation spread outside the contact. Our analysis revealed that the fingertip surface deformation is not confined to the contact interface but extends across the whole fingertip, forming complex strain patterns whose amplitude depends on individual biomechanical and frictional properties. Local curvature changes are also ubiquitous to such loading and might be fundamental to tactile coding (Platkiewicz *et al*., 2016).

## Materials and methods

We adopted an imaging approach called stereo digital image correlation (or 3D-DIC) to quantify fingerpad deformation during controlled mechanical stimulation, a method previously validated in a previous work (Doumont *et al*., 2025). The data analyzed here originates from the same experimental procedures; whereas the prior work focused on a dataset acquired during the initial contact phase, the present work investigates a dataset acquired during the tangential-loading phase.

### Participants

Nine volunteers (two female, 29 ± 5 years of age, mean ± SD) participated in the experiments. They all provided written informed consent and the experiment was approved by the local ethics committee.

### Experimental approach

Briefly, a custom-built platform was used to deliver passive mechanical stimulation to the index fingerpad with a flat transparent surface (glass plate), while the entire fingertip was imaged using multiple synchronized cameras (**Fig.1A, Video M1**). The imaging apparatus consisted of a four-camera array tilted at 30° from the horizontal, forming two stereo pairs (cam#1—cam#4; 2MP, 35-40 px· mm^-1^, 50 fps), and a bottom camera (5MP, 80 px· mm^-1^,100 fps). For the bottom camera only, an alternating light source (at 100 Hz) enabled either uniform illumination of the fingertip or frustrated total internal reflections (FTIR) to capture in-contact regions (**Fig. 1B**). To enable accurate tracking, the fingertip was sprayed to create a fine black-ink speckle pattern. The reconstruction procedure relied on tracking small skin patches (“subsets”) over time that were matched within each stereo pair. Subset coordinates were then triangulated using camera calibration to produce a time-resolved 3D surface of the fingerpad (**Fig. 1C**).

**Figure 1.**
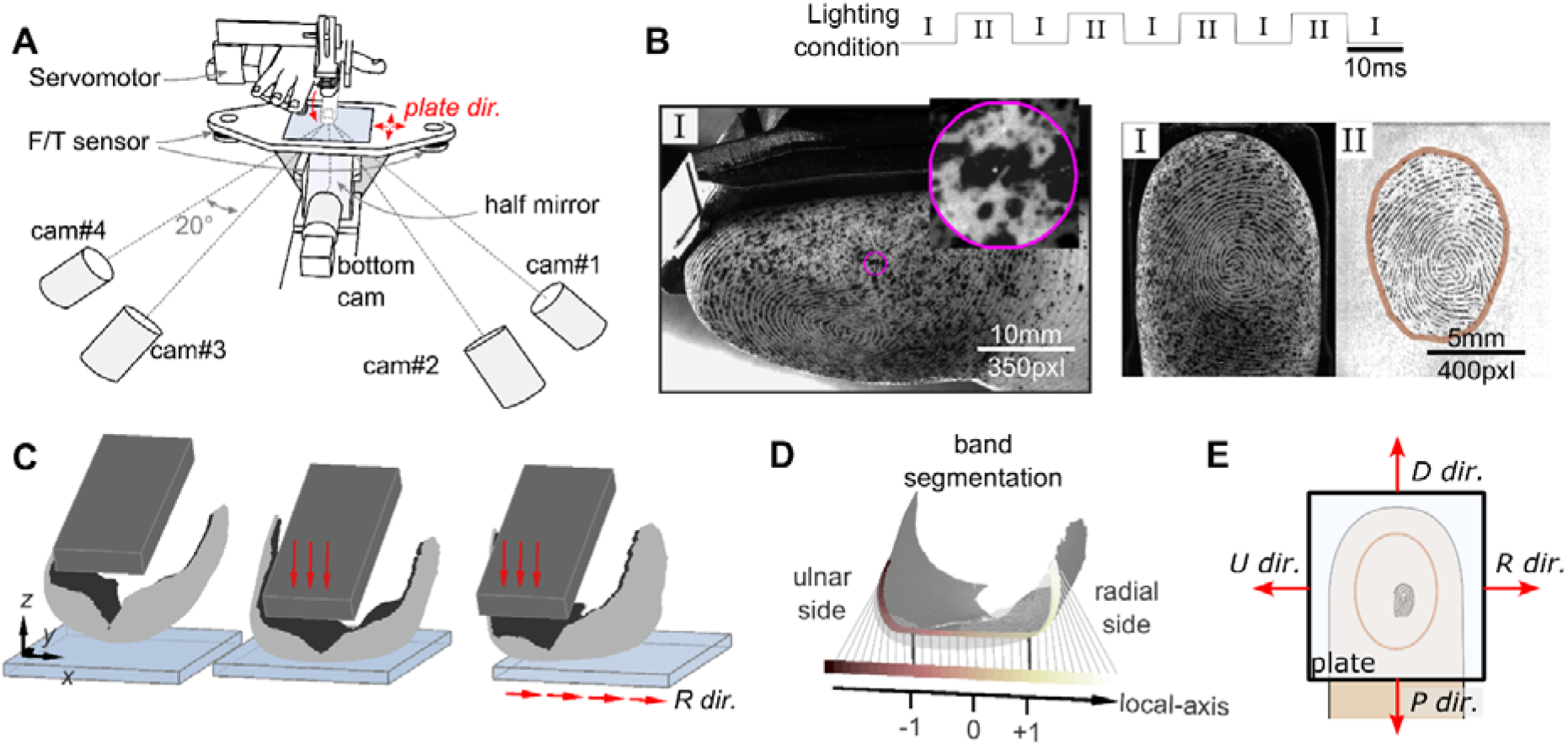
Experimental setup and procedures. (**A**) Schematic of the setup, consisting of a robotic platform and a multi-camera imaging system. Red arrows show the motion of the servomotor guiding the fingertip and the directions of the horizontal motion of the plate. (**B**) Representative raw images of the fingertip acquired using alternating ambient illumination condition (I) or lighting from the bottom (II). Left: image from cam#1 exposed to light condition I (inset: zoomed view of a characteristic speckle pattern). Right: images from the bottom cam acquired under lighting conditions I (middle) to image speckles andII(right) to image fingerprints in contact. (**C**) 3D surface reconstruction of the fingertip at three successive stages: before contact (left), after initial contact (center), and following tangential loading (right) in the radial direction (*R dir*.). (**D**) Illustration of the band segmentation defined along the curved geometry of the fingertip. The band is flattened to create a local axis and normalized to initial contact boundaries (ulnar side boundary = -1; radial side boundary = +1). (**E**) Four plate direction of motion were applied, corresponding to the movement of the plate relative to the nail-fixed fingertip.

### Experimental conditions

After contact initiation, the flat surface was translated horizontally by 15mm to ensure that full slip occurred. Four directions (**Fig. 1E**, ulnar, radial, distal, and proximal directions) were tested at constant speeds of 5 mm/s or 10 mm/s. During this motion, the normal force was maintained at either 1 N or 5 N (the high force/speed condition was not tested). Each trial was repeated four times, yielding a total number of 48 trials per participant (N = 9). Before each trial, the plate was carefully cleaned with alcohol to remove sweat residues. Note that the contact initiation created a non-negligible tangential force in the distal direction. Indeed, because the fingertip was loaded on the plate via rotation around its metacarpophalangeal joint, it pre-constrained the distal region of the fingertip. At a normal force of 5 N, this residual tangential force was 2.17 N ± 0.45 N (median ± 95% CI) in the distal direction.

### Data processing

All data analyses were implemented in MATLAB (MathWorks, Inc. R2021a). Forces and displacements of the robotic platform were sampled at 1kHz and low-pass filtered with a cutoff frequency of 20 Hz. Similarly, displacement and strain fields were filtered with a cutoff frequency of 15 Hz and spatially filtered using a Gaussian kernel of standard deviation 0.2mm. Subsets tracking parameters were adjusted as follows: circular subset radius of 25 pixels and spaced at 5-pixel intervals on the region-of-interest (ROI), yielding an approximate spatial point cloud spacing of 150 μm. The point density of the point cloud was kept for every trial and participant and led to approximately 46000 points tracked over the whole tangential loading. The reconstructed region covered nearly the entire volar surface of the distal phalanx of the index finger (**Fig. 1C**). A narrow unreconstructed region between the two stereo pairs (Doumont *et al*., 2025) introduced an apparent discontinuity that may give the appearance of a crack. A small proportion of trials were lost due to video recording failures: of the 432 trials collected, 416 were successfully recorded and reconstructed (96%). *Gross contact area* — Tracked points were considered in contact if their z-coordinate fell below a neighborhood of the contact fitted plane. The gross contact contour area was defined as the boundaries of the contacting points. The gross contact area will be referred to as the ‘contact area’ in the rest of the paper (example shown with a brown contour in **Fig. 1B**).

#### Slip/stuck area

In-contact tracked points were classified as slipping if their speed fell below 95% of the plate velocity; conversely, in contact-points whose velocity remained above this threshold were classified as stuck, hence moving with the plate.

#### Friction

The static coefficient of friction (*CF*_*stat*_) was defined as the ratio between the tangential force and the normal force at the instant of full slip, i.e. at the moment when the stuck area becomes null.

#### Band region segmentation

To characterize the spatial evolution of deformation inside and outside the contact, a 2-mm wide band across the fingertip was defined along the ulnar-radial axis, centered on the contact area. The band was further divided into 1-mm bins to quantify local deformation along the curved geometry of the fingertip. Bin locations were then normalized to the contact boundaries to allow for comparisons across trials and participants (**Fig.1D**).

#### Surface strain

Surface strain field was computed from the deformation gradient tensor derived from the surface-triangulated point cloud maps using the MultiDIC toolbox (Solav *et al*., 2018). The reference fingerpad state was taken at the end of the initial contacting phase, such that only the strains associated with the tangential loading phase are reported.

#### Geometric curvature

To quantify changes in the global geometrical shape of the fingerpad, we computed the surface curvature for each point-cloud map. This metric also referred to as *surface variance*, examines how much the local point cloud varies in the direction of the estimated normal (Bae & Lichti, 2008). It was computed from the eigenvalues of the covariance matrix of point coordinates within a local neighborhood, with curvature defined as the smallest eigenvalue normalized by the sum of all eigenvalues. Larger values indicate greater deviation from a locally planar surface and therefore exhibit higher curvature. Because both point-cloud resolution and the number of neighbors used in the surface curvature estimation were fixed, this metric is suitable for comparisons across trials and participants.

#### Strain energy

As the plate translates horizontally, the fingertip resists the motion, and a part of the stimulus mechanical energy is transferred to the fingertip and stored as deformation in the skin. Assuming an isotropic material and hyperelasticity, appropriate for large deformation (>10%), it is possible to estimate the energy density associated to the surface strain measured here. Using material properties in the range of *in-vivo* measurements (Young’s modulus E = 1MPa; Srinivasan *et al*. (1992)), and assuming volume incompressibility (ν = 0.5), the strain energy density u (mJ.mm^-3^) can be expressed in terms of Green-Lagrange principal strain *E*_*i*_ using a Neo-Hookean formulation of the energy (**Eq. 1**). The normal component to the surface *E*_3_—not being measured here—is computed as a combination of *E*_1_ and *E*_2_ under the incompressibility assumption J (**Eq. 2**).

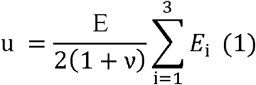

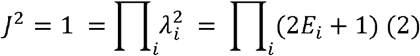

The total strain energy U (mJ) was obtained by integrating the strain energy density over a given volume (Delhaye *et al*., 2016): As we do not measure bulk strains – inside the tissues, we made the simplifying assumption that strains are uniform across depth, with skin depth affected by strain estimated at 2mm. This is supported by in-bulk measurement showing that skin remains relatively homogeneous perpendicular to the surface under similar loading conditions (Corniani *et al*., 2025). The total energy is thus approximated by summing the contributions of each triangular surface element of area A_i_ over the reconstructed surface S (**Eq. 3**).

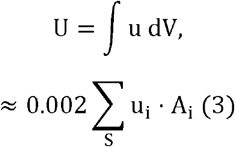

#### Fingertip stiffness

A proxy of the fingertip stiffness of each participant was evaluated from the load-displacement characteristic of the contact initiation phase. As the displacement of the fingertip support was not measured, the indentation ξ was approximated as the relative motion between points identified as being in contact (i.e. approximately coplanar) and points near the nail, which were minimally affected by the applied load (i.e. the normal force during contact initiation). Assuming Hertz contact power law *NF* = *a* ξ^3/2^, we fitted the load-displacement curve of each participant in the force range [0,1N]. The parameter *a* was then used as a proxy for each participant’s fingertip stiffness.

#### Fingertip width

We measured the fingertip width as a proxy of the participant-specific fingertip size. This was a reasonable assumption given that contact area was shown to scale well with fingertip width over the same population as this study. Fingertip width was evaluated once, using the bottom camera at a frame just before contact, when the fingertip was in undeformed state.

### Statistical analysis

We used linear mixed-effect models (LMMs) to examine the factors affecting strain energy density. Model 1 tested the effect of experimental conditions, Model 2 assessed the impact of the frictional properties of the contact, and Model 3 evaluated the influence of individual fingertip stiffness and size. Each model was structured differently to address the specific predictors of interest, allowing appropriate inference at the corresponding level—within or between individuals—while accounting for potential confounding effects, such as the influence of experimental conditions.

#### Model 1

We tested the effect of the experimental conditions on the median energy density (u), allowing for between participants (SUBJ) variability to be accounted for in terms of model intercept. Interaction terms were kept when supported by experimental design (high normal force combined with high speed was not evaluated) and were further explored when significant with post-hoc analysis (**Eq. 4**).

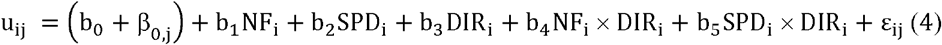

where *u*_ij_ is here the median energy density of the i-th trial associated to the j-th participant, b_0_ to b_5_ are the fixed effects related to normal force condition (NF), plate speed (SPD) and plate direction (DIR) with possible interaction effects. The parameter β_0,j_ ∼N(0,σ_0_^2^) is the random intercept associated with the factor SUBJ, and ε_ij_ ∼ N(0,σ^2^) are residuals of the model.

#### Model 2

To evaluate the effect of the frictional properties of the contact on intra-participant variability in stain, the median strain energy density was predicted from the static coefficient of friction (CF). It captures differences in friction-energy coupling specific to each participant in each condition context, treated as random intercepts (**Eq. 5**).

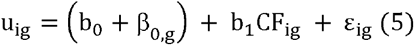

where indices are as in Model 1, g(j,k) is the grouping factor associating a participant and the k-th combination of experimental conditions (combination of NF, SPD and DIR conditions). The number of combinations was further reduced if an experimental condition was found non-significant in *Model 1*. The parameters b_0_ and b_1_ are the fixed intercept and slope, and β_0,g_ ∼N(0,σ_0_^2^) is the subject-condition specific random intercept, and ε_ig_ ∼N(0,σ^2^) are residuals.

#### Model 3

To assess whether fingertip geometry (Width) and fingertip stiffness (Stiff) account for inter-subject variability in the strain energy, this model treated only experimental condition combinations as random intercepts, in contrast to Model 2 (**Eq. 6**).

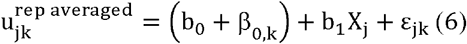

where 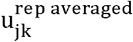 is the averaged median strain energy density across repetitions of the j-th participant, associated with the k-th experimental condition combination. The averaged value was taken to remove imbalanced number of trials between participants due to lost of data (see *Data Processing* section). b_1_ is the fixed effect for the independent variable X_j_, either Width or Stiff. The parameter β_0,k_ ∼ N(0,σ_0_^2^) is the condition specific random intercept associated with condition combination k, and ε _jk_ ∼ N(0,σ^2^) are residuals.

## Results

We reconstructed the 3D shape and deformation fields of the fingerpad surface while it was normally loaded (see Doumont *et al*., (2025)) and slid tangentially against a flat plate. First, we validated the measurements made with stereo DIC (3D) against independent measurements made in 2D at higher resolution, which have been previously used and validated (Delhaye *et al*., 2014, 2016; de Dunilac *et al*., 2023). Both measurements of the strain fields were compared within the contact area during tangential loading (**Fig. 2A**). We found a robust point-wise match between the two observations (**Fig. 2B**). The analysis demonstrates high precision, evidenced by a high intraclass Correlation Coefficients (ICC>0.9) indicating low random noise, and a strong linear association (r = 0.94). In addition, Deming regression slopes for each strain component were close to unity (ranging from 0.81 to 1.17), indicating minimal systematic bias between methods.

**Figure 2.**
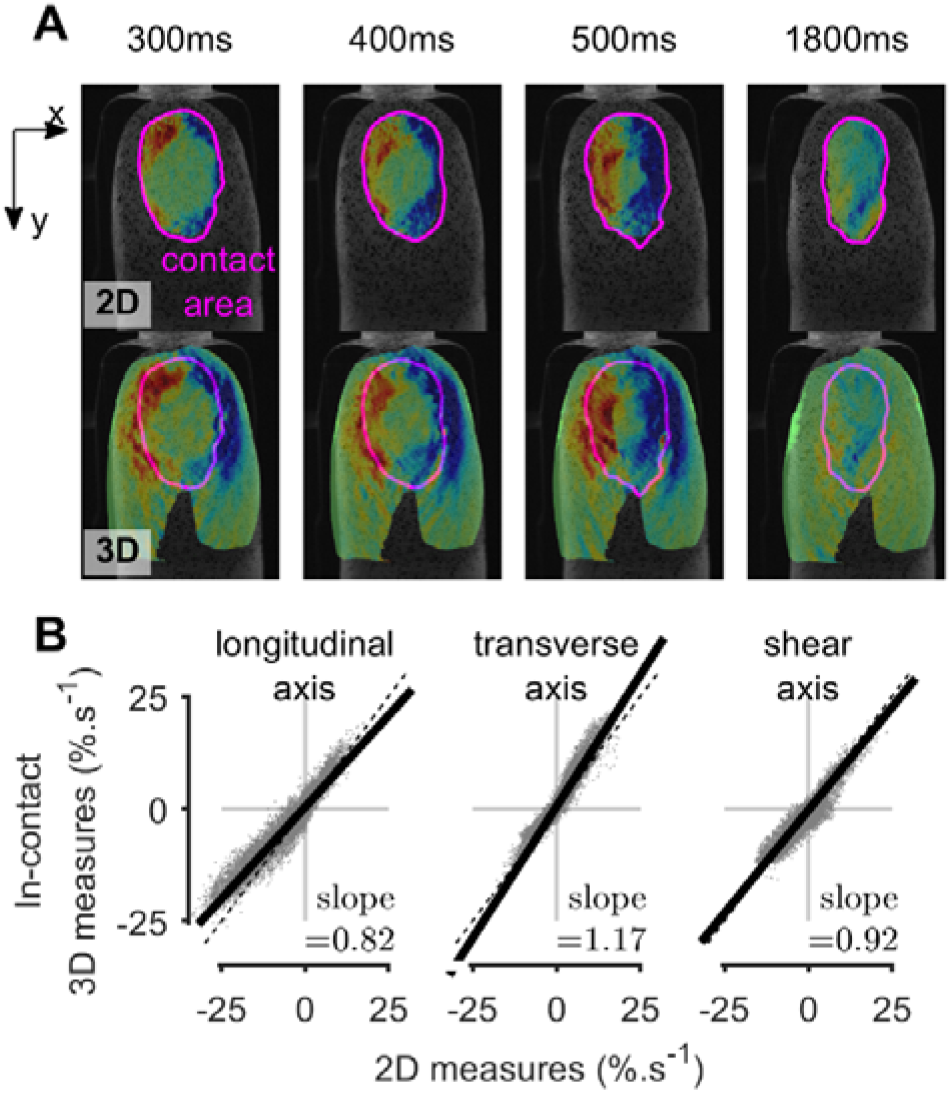
Validation of 3D measurements using simultaneously recorded in-contact 2D data for a representative trial. (**A**) Spatiotemporal strain rate fields, shown for the component aligned with plate motion axis (here, positive x). Measurement from the 2D bottom camera (top row) are compared from the 3D stereo-measurement (bottom row), projected onto the bottom-camera images. The 3D measurement coordinates were mapped into the bottom-camera reference frame. Projected coordinates aligned with the physical fingertip geometry seen from bottom-camera images. (**B**) Global point-wise comparison across the full spatiotemporal continuum. All data points recorded over the contact area (co-registered onto a common grid of 0.3mm spacing) and throughout the tangential-loading phase are pooled to assess the agreement of the two measurement methods. Comparisons of strain rates were done for each strain component, i.e. in the longitudinal, transverse and shear axis. Thick black lines indicate linear fit; perfect agreement would correspond to a unit-slope (dashed black line) passing through the origin.

### Whole fingertip deformation during tangential loading

Loading the index fingerpad tangentially led to widespread deformation across the entire fingertip, both locally and globally, regardless of loading direction. **Figure 3** illustrates this behavior for the ulnar direction, a stimulation direction predominantly seen during object manipulation due to the gravity force vector (see **Figs. A1-3** for similar plot in other experimental directions, and **Video M2**). As the plate translated laterally, frictional forces at the skin-glass interface progressively pulled the skin in the plate direction (**Fig. 3A**). Thus, the last stuck region of the contact follows the plate motion up until full slip occurred. Consistent with previous observations, partial slip developed progressively within the contact area; the transition from a fully stuck to a fully slipping state initiated at the periphery and propagated inward to the center of the contact (**Fig. 3B**, see location of the last stuck region). Importantly, the evolution of surface strains closely followed this slip phenomenon and extended far beyond the contact area (**Fig. 3C, E, G**). Regions outside of contact – unstuck and therefore free to move – were the first to deform, forming a wavefront that propagated inward within the contact area as the stuck region shrank. As a result, deformation was largest near the contact, whereas the last stuck, central, region presented minimal strain.

**Figure 3.**
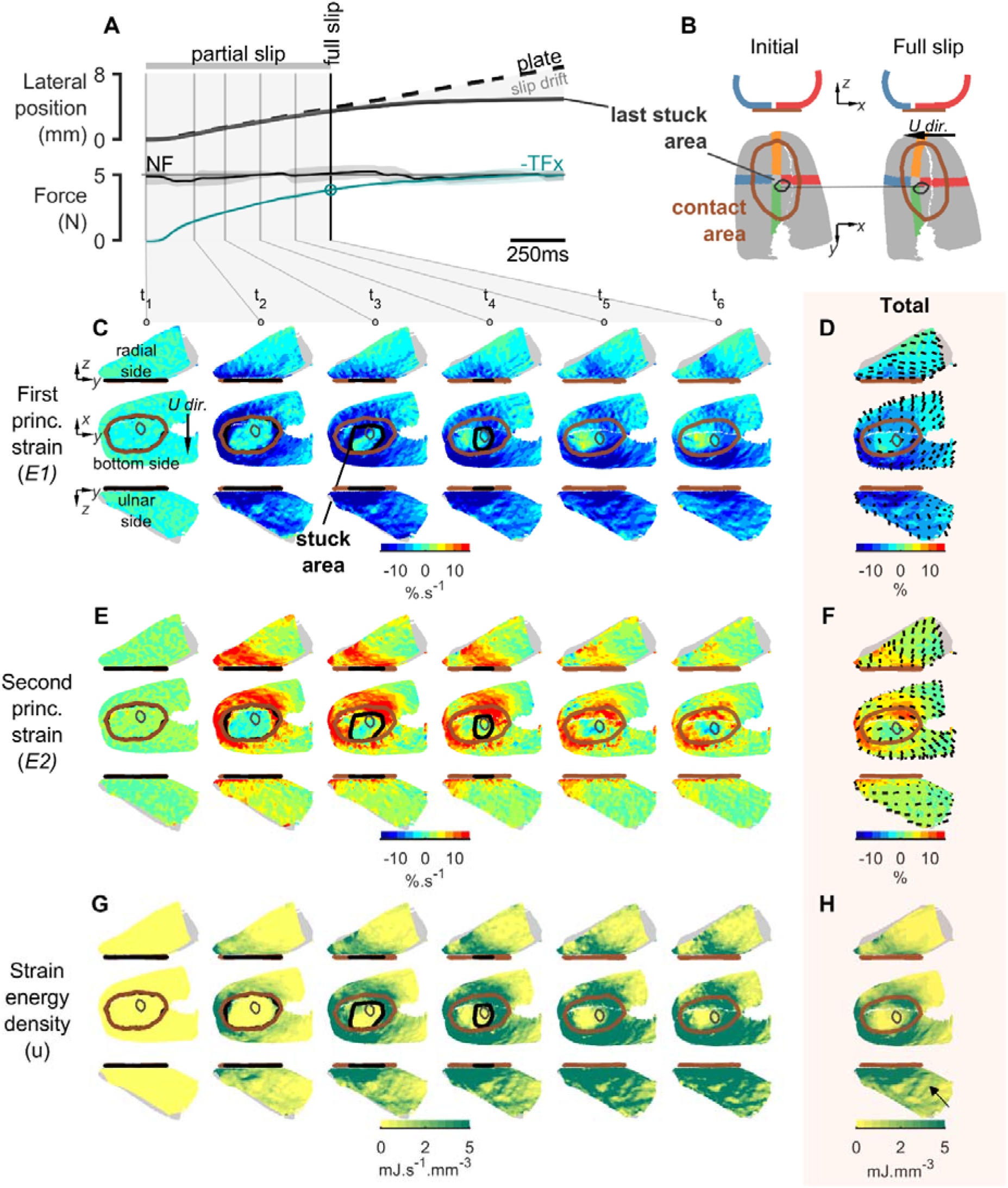
Typical whole fingertip deformation encountered during tangential (ulnar) loading (S06). (**A**) Evolution of the global variables as a function of time. Top: Lateral position of the plate and of the last part of the skin stuck to the plate. Once the full slip is reached, relative position drift is observed. Bottom: Controlled normal force (black) and tangential force produced in the direction of stimulation (cyan). Solid lines show mean values across repetitions of the same condition, with shaded areas indicating ±1 SD. (**B**) 3D maps of the fingertip reconstruction at initial contact and at full slip. The location of the last stuck region is indicated with crossed bands to help visualize the gross deformation. (**C**) 3D heatmaps of the strain rate evolution during partial slip of the first principal component (E1) at time points defined in (A). Views are shown from the radial side (top), the bottom side (middle), and from ulnar side (bottom). Note that the opposite-facing side is colored in gray. The boundary of the contact area is overlaid in solid brown, the current stuck region in solid black, and the last stuck region is shown in gray. (**D**) 3D heatmaps of total strain (integral of strain rates) for E1, shown in the same perspectives as in (C). Local directions of E1 are overlaid. (**E-F**) Same as (C-D), for the second principal strain component (E2). (**G-H**) Same as in (C-D), for the strain energy density.

The strain rate amplitudes generated during the controlled linear motion were comparable to those reported during manipulation (grip and lift tasks), which typically peak at approximately 20%.s^-1^ (Schiltz *et al*., 2022; Delhaye *et al*., 2024). Larger surface contractions amplitudes were observed on the ulnar side of the fingertip (99^th^ percentile of total strain of about 60%; **Fig. 3C-D**), whereas the radial side tended to exhibit larger stretches (**Fig. 3E-F**). Local strain orientations were broadly aligned with the direction of the plate motion while conforming to the curved geometry of the fingertip (see local strain orientations overlaid on the total strain maps).

Looking at the stored elastic energy density provided a convenient scalar that gives an intuition of the repartition of the deformation intensity (**Fig. 3G-H**). On the side of stimulation and outside the contact boundaries, elastic energy increased proportionally with tangential loading. Moreover, we observed an asymmetrical distribution of the total energy density when comparing the ulnar and radial sides of the fingertip (top versus middle row). In addition, non-uniform localized concentration patterns of strain energy were observed (**Fig. 3H**; black arrow and **Fig. 4A-B**). These complex, curvilinear patterns–occurring in regions of contraction—could result from skin wrinkling induced by tangential loading.

**Figure 4.**
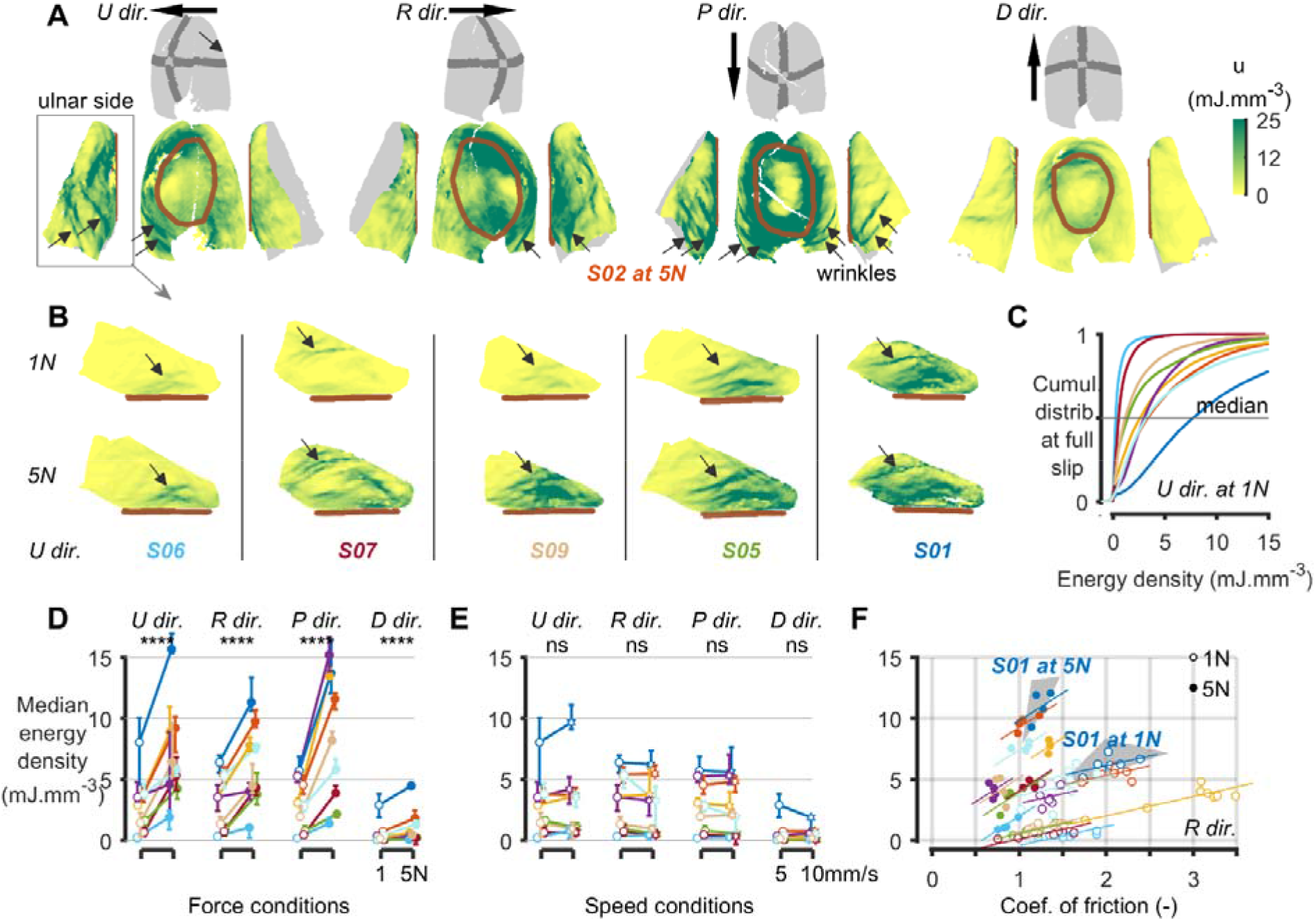
Influence of loading conditions on the deformation patterns and intensity. (**A-B**) Maps of total strain energy density at full slip; (A) in all plate directions (U,R,P,D dir.) for a single participant (same view as in Fig.3), and (B) in the two normal force conditions (1N and 5N) across five participants in the ulnar direction at full slip (ulnar side view). Black arrows indicate wrinkling, visible as local longitudinal concentration of energy density on the side of the stimulation direction. Individuals are sorted by increasing energy. (**C**) Cumulative distribution of the energy density over the entire fingertip reconstruction at full slip for all participants, highlighting high variability in energy density distribution across participants. (**D-E**) Effect of (D) force conditions and of (E) speed conditions on the median energy density for each plate direction for all participants. The error bar shows the standard deviation over repetition of the same condition. (**F**) Effect of the static coefficient of friction on the median energy density for all participants at 1N and 5N in the radial plate direction. Each color corresponds to a specific participant in the figure.

### Effect of loading direction, force, speed, and participant variability on fingertip deformations

As expected, strain energy patterns appeared stereotyped to the loading direction of the fingerpad (**Fig. 4A**) while loading the fingerpad at higher normal force predominantly impacted the deformation amplitude, not its spatial distribution (**Fig. 4B**). Remarkably, we observed similar wrinkles for all plate directions across all tested experimental conditions, and all participants (see black arrows in **Fig. 4A-B**; **Video M3**). While wrinkles varied in patterns depending on participant and stimulation direction some qualitative trends were observable. First, wrinkles were appearing in regions undergoing contraction. Second, they extended perpendicular to the principal direction of contraction. Third, they were predominantly seen in proximal regions of the fingertip, where the soft pulp of the tissue is thicker.

To quantify the global effects influencing the amplitude of strain energy, we summarized the distribution of total strain energy density at full slip using its median value (**Fig. 4C**), which provided a scalar descriptor of deformation intensity while limiting the influence of localized extrema.

Regarding the effect of experimental conditions on strain energy (see Methods, Model 1), interaction effects were found between normal force (NF) and plate direction (DIR) (p < 10^-4^), and between plate speed (SPD) and direction (p < 10^-4^). Post-hoc analyses revealed a strong positive effect of normal force in all four plate directions (**Fig. 4D**; mean ratio = 3.2; p < 10^-4^) whereas speed did not affect the energy median density in any direction (**Fig. 4E**; p > 0.05). Thus, the speed only affected the rate of deformation and not the total amplitude of strain. Then, we tested whether the plate direction affected strain energy. No clear effect of direction was detected in the ulnar-radial axis (p = 0.14). In contrast, a marked effect of direction was found along the proximal-distal axis: the median energy density in the distal direction was approximately one quarter of that measured in the proximal direction (mean ratio = 0.26; p < 10^-4^). This asymmetry was not fully explained by the direction-dependent variation in tangential force, which differed by a factor of about two between proximal and distal stimuli (see *Methods*). It could reflect intrinsic anatomical and geometrical factors: the shape of the distal phalanx and the relative angle between the fingertip and the plate may lead to asymmetric distribution of contact pressures. The reduced amount of soft tissue at the distal end of the fingertip could further limit its capacity to deform, thereby contributing to the observed imbalance in energy density (Johansson & Flanagan, 2009).

To assess whether the gross spatial distribution of deformation depends on motion direction, we compared the median energy density at full slip between the ulnar and radial lateral sides (outside the contact) of the fingertip. Consistent with observations in **Fig. 3G** (first vs last row), we observed a strong reorganization of the strain energy magnitude distribution depending on plate direction (p < 10^-4^). Specifically, strain energy was higher on the side facing the motion direction, indicating that the compressed side experienced greater strain energy.

We reported both intra-individual variabilities (across repetitions of the same conditions), and large inter-individual variability (across participants under identical conditions). First, we examined whether the frictional properties of the contact could influence the intra-individual variability in fingertip deformation (**Fig. 4F**, see also *Methods*, Model 2). We exploited the natural trial-to-trial variations in the static coefficient of friction across repetitions (mean standard deviation of 0.20 at 1 N and 0.11 at 5 N). These changes could be due to occlusion, sweat rates or sweat residues from previous trials, as moisture levels are known to fluctuate significantly during object manipulation (André *et al*., 2010; Barrea *et al*., 2016). Accounting for experimental conditions using random intercepts, the model revealed a positive effect (p < 10^-4^; marginal R^2^_adj_ = 0.97), indicating that friction increased deformation. This indicates that friction is the dominant driver of trial-to-trial deformation variability, indicating that variations in interfacial friction strongly modulated fingertip deformations for each individual.

Finally, we investigated the potential sources of the inter-individual variability (see *Method*, Model 3). We tested whether fingertip stiffness and width, affected deformation across participants. When considered independently, the *stiffness* explained a large fraction of the variance at the population level (p < 10^-4^; marginal R^2^_adj_ = 0.51), with stiffer fingertips exhibiting lower deformation. *Width* alone also showed a significant, but weaker negative effect (p=0.033; marginal R^2^_adj_ = 0.42). Though, these predictors are not independent: the measured proxy for fingertip stiffness reflected an effective mechanical response that depends on both the tissue material properties and fingertip geometry. Consistent with this interpretation, while adding *width* as an independent predictor in the *stiffness*-only model does not improve the fit (p = 0.54; ΔAIC = 1.63; ΔBIC = 4.32, marginal R^2^_adj_ = 0.50), allowing for interaction effect between the two predictors resulted in a substantial improvement of the model (p < 10^-4^; ΔAIC = -49.55; ΔBIC = -44.17) with marked increase variance explained by the fixed effects (marginal R^2^_adj_ = 0.72).

Overall, these results indicated that intra-individual variability in the amplitude of fingertip deformation was largely explained by variations in friction, whereas inter-individual differences were well accounted for by fingertip size and stiffness considered jointly.

### Out-of-contact surface deformation overwhelms within-contact deformation

To understand the gross distribution of fingertip deformation and its evolution during tangential loading, the elastic energy stored within-contact was separated from outside-contact. We showed that both within- and out-contact power (energy rate) were increasing and peaked near full slip (**Fig. 5A**). The behavior was consistent across all participants regardless of the distance to full slip. The out-of-contact energy rate peaked significantly earlier than within-contact (estimate = -16.4%; p < 10^-4^), occurring at 76.49 ± 10.1 % (mean ± SD) of the distance of full slip, compared with 91.7 ± 8.9 % for the within-contact energy. Consistent with previous observations (Delhaye *et al*., 2014), residual deformation took place after full slip. This can be explained by tangential force still increasing after full slip (**Fig. 3A**), and by possible release of residual constraints inside the contact region due to the normal loading phase.

**Figure 5.**
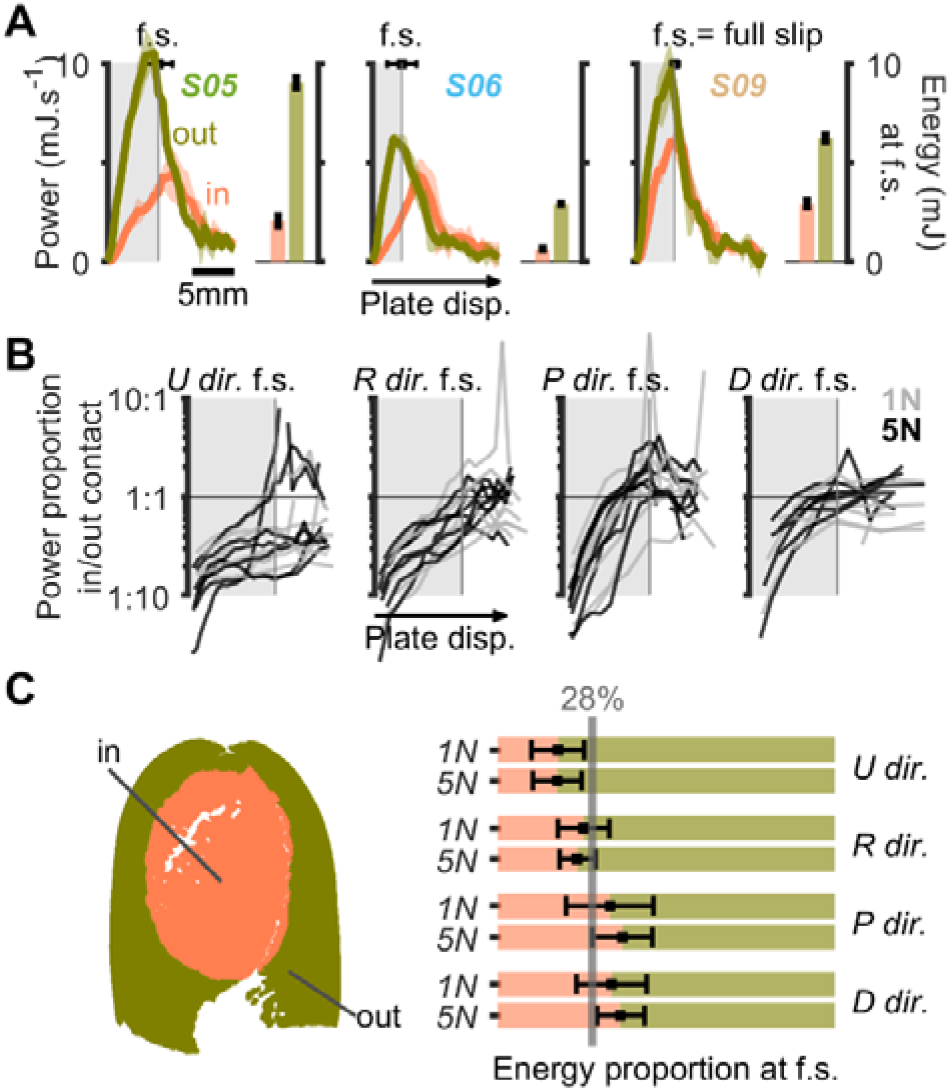
Quantitative comparison of outside vs inside contact deformation energy. (**A**) Evolution of strain energy rate is shown as a function of plate displacement (left subplots) and total strain energy at full slip (right subplots), for three participants in the radial direction at 5N. Mean values and standard deviation; the timing of full slip is indicated with its variability. (**B**) Evolution of the power proportion as function of the plate displacement normalized to full slip distance. Evolution traces are shown as the average over trial repetition for all participants, directions (U,R,P,D dir.) and normal forces conditions (1N in grey and 5N in black). (**C**) Proportion of total strain energy versus normal force and plate directions are shown for all participants. Mean values and standard deviation are shown. Less than a third of the total strain energy is stored inside the contact.

The temporal progress of deformation rate from outside to inside-contact as seen in **Fig. 3** was also reflected in the evolution of their relative contribution to the total deformation power (**Fig. 5B**). At the onset of tangential loading, deformation power was strongly dominated by out-of-contact regions. As partial slip increased, the contribution of in-contact deformation progressively increased.

More strikingly, a strong imbalance in energy distribution was observed at full slip. The energy accumulated outside of the contact region was two to three times higher than that stored within the contact area (**Fig. 5C**). It showed that deformation was not at all constrained to the contact area but spread markedly outside of it.

Finally, we found that part of variance in the proportion of energy stored inside the contact was depending on the fingertip size and stiffness (R^2^ = 0.45; same statistical analysis as model 3). That is, higher proportion of deformation inside the contact with larger and/or stiffer fingertips (p < 10^-4^). This suggests that the deformation spread during tangential loading is influenced by the fingertip size and stiffness.

### Smooth transition between in and out of contact deformation

To study the temporal evolution of strain and the spatial continuity of deformation between skin regions inside and outside the contact, we analyzed in detail the deformation along a linear band crossing the fingertip from radial to ulnar and passing by the center of contact (see Methods). The band was flattened (**Fig. 1D**), and strains were expressed in a coordinate system aligned with the band axis. Strain rates were developing first outside the contact and progressively propagated inward as partial slip occurred (**Fig. 6A)**. After full slip, the total strains were largest near the contact boundary, but there was a very smooth continuity of the deformation from inside out (**Fig. 6B**). The fingertip experienced contractive strain (negative) on the side in the direction of motion (leading side) and tensile strain (positive) on the opposite side (trailing side). A notable contractive strain drop just outside of the contact was observed, consistent with instabilities created by wrinkling of the skin. Nearly mirrored profiles of total strains were observed for ulnar and radial plate directions.

**Figure 6.**
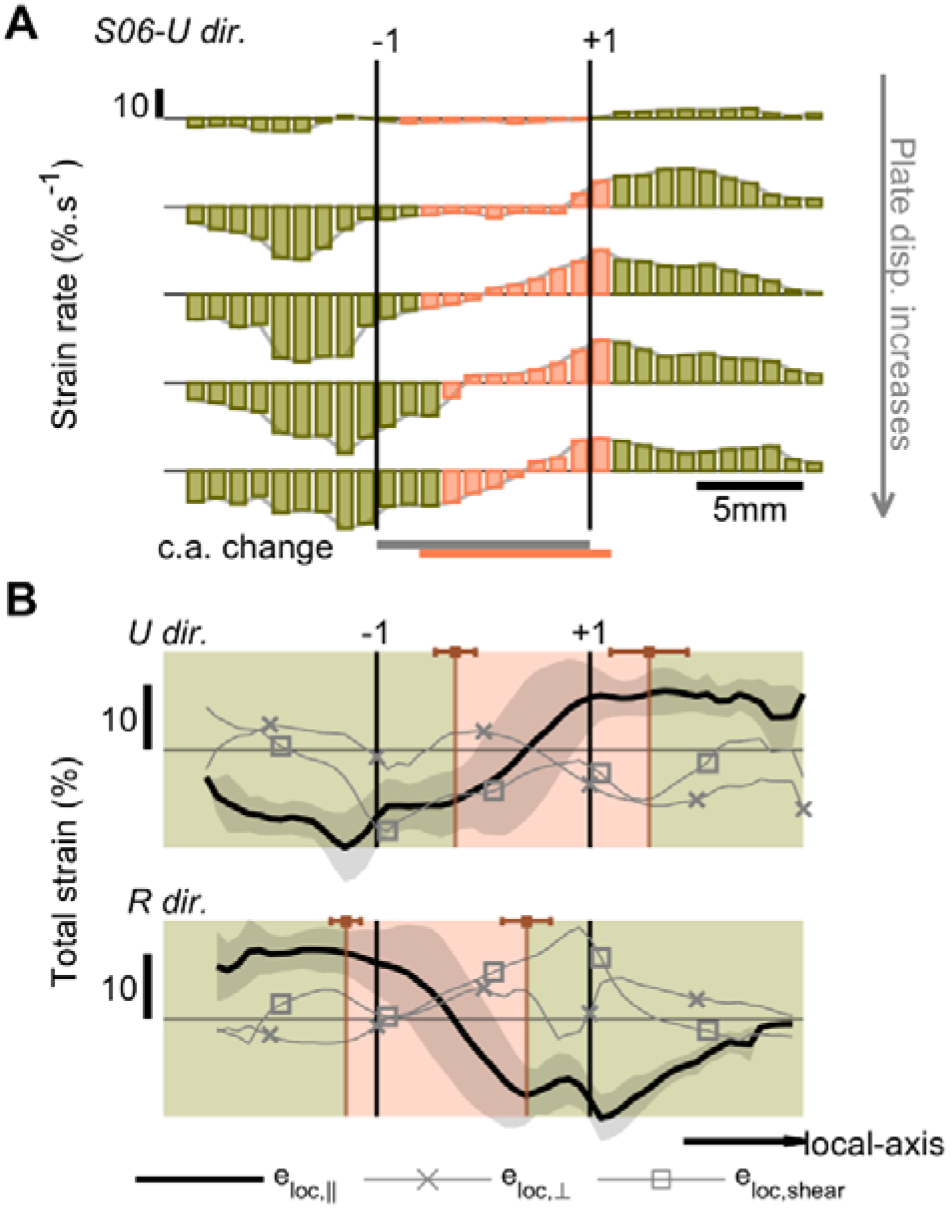
Continuity of the deformation outside vs inside the contact. (**A**) Strain rate profiles along the band region and their evolution as partial slip increases (top to bottom), shown for an illustrative trial. The strain component shown is expressed in the local coordinate system, aligned to the plate motion. (**B**) Total strain profiles along the band region for ulnar (*U dir*.) and radial (*R dir*.) plate directions at full slip. Each strain component is shown, with the component aligned to the local-axis (e_loc,||_) plotted in solid black line and standard deviation in shaded area. Grey line with crossed markers for the strain perpendicular (e_loc, ⊥_) and with squared markers for the shear strain (e_loc,shear_) to the band axis. The region inside the contact area is shown in orange and outside the contact in green.

In addition, this analysis showed that the contact area shifted along the fingertip surface in the direction opposite to the plate motion, consistent with a rolling motion of the fingertip during tangential loading.

### Sharp changes in finger curvature during loading

So far, the analyses have focused on surface skin deformations. Whereas it is difficult to interpret the measurement in terms of the bulk deformation as we cannot measure inside the tissues, the changes in curvatures provide a hint at the deformation in the inner tissues. We measured the skin curvature in and close to the contact (**Fig. 7A**). The fingertip skin was highly curved around the contact boundaries, as expected after initial contact against a flat rigid plate. Surprisingly, we observed large changes in local curvature during tangential loading (**Fig. 7A**, lower panel).

**Figure 7.**
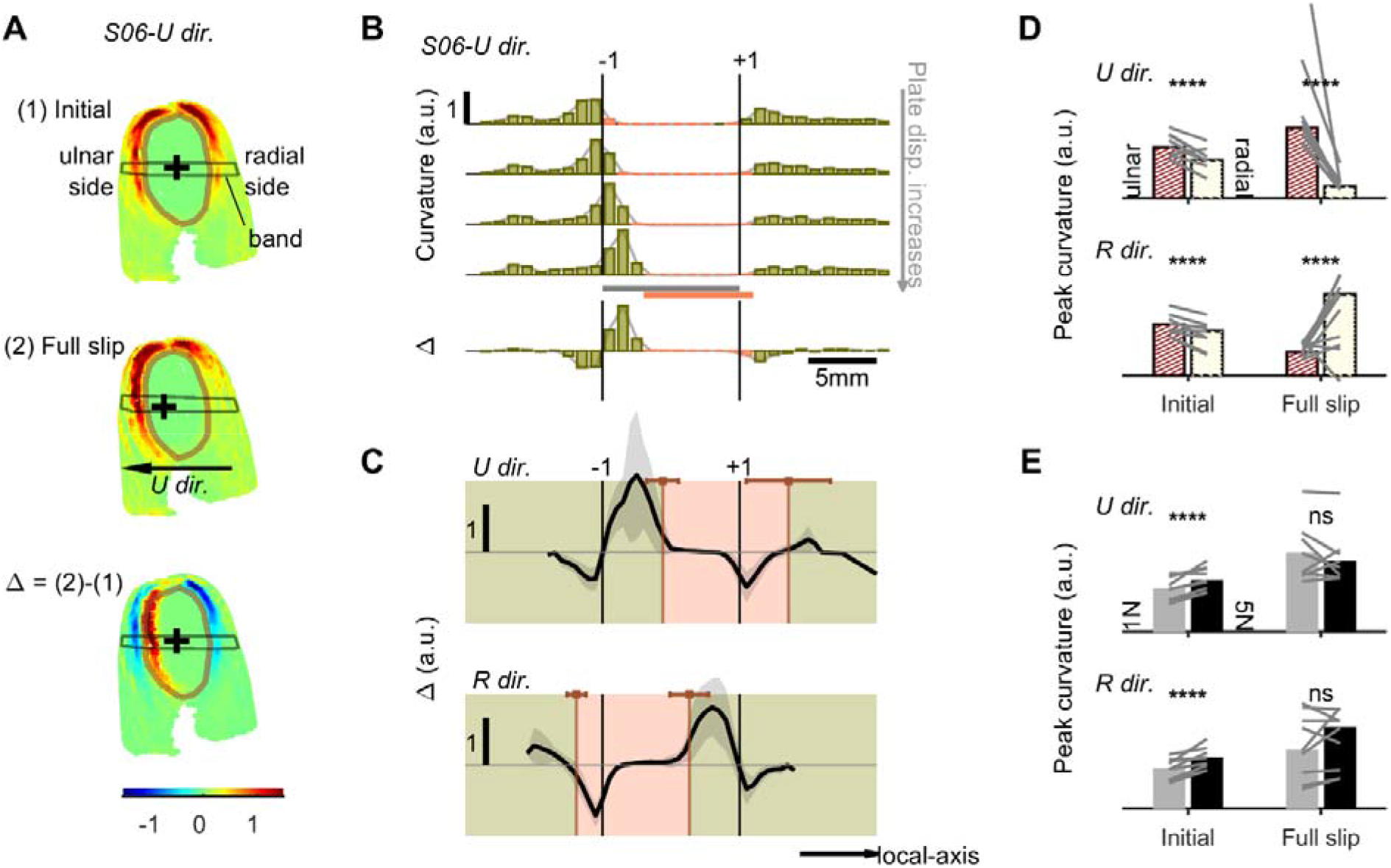
Skin surface curvature. (**A**) Map of local curvature for an illustrative trial: fingertip initially loaded against the plate (top), after plate motion (middle), and the resulting curvature change expressed in the initial configuration (bottom). The black cross tracks a portion of the skin and illustrates the fingertip motion. (**B**) Local curvature profiles along the band region plate displacement increases for the same trial as in (A); the last row shows the total curvature change. (**C**) Total change of curvature along the band region for opposite plate directions (top: ulnar; bottom: radial), for all participants at full slip. The region inside the contact area is shown in orange and outside the contact in green. (**D**-**E**) Peak curvature before and after plate motion for opposite direction at full slip (top: ulnar direction; bottom: radial direction). Peak curvature was quantified on each side of the fingertip in (D), and across the entire fingertip for the two normal force conditions in (E). Statistical significance is indicated as ns (non-significant) and **** (p < 10^-4^).

Looking again along the same radial-ulnar band as in **Fig. 6**, as tangential loading increased, the location of the peak curvature progressively shifted along fingertip, reflecting rolling of the fingertip over the plate (**Fig. 7B**). Consequently, skin regions that were initially flat (in contact with the plate), were later pushed outside the contact area and experienced pronounced changes in curvature (**Fig. 7C**; see location of the peak of the changes in curvature). Interestingly the location of the peak change of curvature aligned well with the sharp drop in contractive strain reported in **Fig. 6C**.

Then, we assessed whether curvature was symmetric across the two sides of the fingertip. Peak curvature was already asymmetric between the ulnar and radial sides at initial contact (ulnar/radial ratio = 1.38; p < 10^−^□), indicating that fingertip geometry was not perfectly symmetric immediately following contact onset (**Fig. 7D**). This asymmetry likely reflects the natural pronation of the fingertip toward the thumb when the finger is extended (Bansal & Craigen, 2007). The nail of the fingertip was thus tilted, introducing a geometric asymmetry. After tangential loading, the curvature imbalance became much more pronounced (ratio = 5.13; p < 10^−^□) and depended on plate direction: for ulnar-directed stimuli, curvature was larger on the ulnar side than on the radial side.

Finally, we examined the effects of experimental conditions and individual biomechanical properties on the peak curvature along the band. Peak curvature expectedly increased with normal force at initial contact (**Fig. 7E**; estimate = +23.1% between 1N and 5N conditions; p < 10^−^□). In contrast, this effect disappeared after tangential loading (p = 0.95). In addition, peak curvature depended on individual fingertip width (p < 10^−^□), with smaller fingertips exhibiting higher curvature for the same applied normal force. This effect of fingertip width was present both at initial contact and after tangential loading. *Stiffness* also had a significant negative effect on peak curvature at initial contact (p = 0.0090) but was non-significant after tangential loading (p = 0.058).

## Discussion

This study quantified how the entire fingertip surface deforms under tangential loading, revealing significant and widespread strains that extended well beyond the contact region. Rather than being confined to the interface, deformation consistently originated outside the contact and propagated inward as the interface transitioned from full stick to full slip. Tangential loading also generated pronounced directional asymmetries, local compressive instabilities consistent with skin wrinkling, and substantial changes in local curvature, indicating both surface strains and bulk tissue deformation.

### Limitations

The present study used a flat rigid contact surface, which differs from many natural interactions where objects often present curved or irregular geometries and may be compliant. Such geometries would likely modify the distribution of contact pressures and the resulting skin deformation patterns (Srinivasan & Dandekar, 1996; Xu *et al*., 2021). Nevertheless, the simplified configuration allowed us to precisely quantify the evolution of surface strains across the entire fingertip and isolate the mechanical consequences of tangential loading. While strain-rate levels were compatible with object manipulation (i.e. in the range of 30%/s for grip and lift, Delhaye *et al*., 2021*b*), force dynamics during natural interactions are typically more complex, involving time varying normal forces and sliding speeds (Forssberg *et al*., 1991; Gueorguiev *et al*., 2022). Finally, the precision of the reconstruction could be further improved by incorporating additional stereopairs of cameras to better account for the fingertip geometry, thereby reducing geometric image distortion (Doumont *et al*., 2025).

### Implications for manipulation and surface exploration

From a manipulation perspective, partial slip creating strain wave within the contact is thought to inform the nervous system about contact stability (Johansson & Westling, 1984; Schiltz *et al*., 2022; Delhaye *et al*., 2024). Our results suggest that out-of-contact regions could provide additional information about early mechanical events associated with partial slips inside the contact as deformation started outside the contact. This early signal could potentially provide rapid adjustments of prehension forces together with proprioceptive inputs.

This region continued to consistently deform for the whole tangential loading and strain energy rate peaked around full slip. One can reasonably assume that under high-friction conditions, at the skin surface interface, the outside region would constitute the only region free to deform, while the contact remains stuck and mechanically fully constrained. Such a spatial segregation of deformation could therefore provide distinct sensory cues: out-of-contact deformation reflecting tangential force build-up, and in-contact deformation signaling slip onset from the strain wave propagation. This hypothesis could be tested in future work, using for instance stimuli of distinct frictional properties.

Additionally, we showed that magnitude of the deformation scaled with normal force and was influenced by stiffness and size of the fingertip. In active condition when exploring surfaces, it was shown that individuals with stiffer skin and larger fingertips tend to apply a larger normal force (Kurimoto *et al*., 2026). One hypothesis would be that individuals regulate their applied force to achieve comparable levels of skin deformation, probably affecting tactile perceptual discrimination (Li & Gerling, 2023).

### Consequence in terms of neural coding

Studies with neurophysiological recordings have consistently reported that virtually all afferents innervating the fingertip actively respond during tangential loading (Birznieks *et al*., 2001; Jenmalm *et al*., 2003; Wheat *et al*., 2010). The strain patterns here provide a mechanical basis for this widespread neural activation: substantial strain energy was observed across the whole fingertip, with energy outside the contact exceeding that within the contact by a factor of two to three after full slip. This shows that population activity could even be dominated by non-contact skin deformation, notwithstanding the responses related to the deformation normal to the surface that weren’t measured in the current study.

Previous studies have shown that spike timing of FA-I afferent were particularly efficient at discriminating quickly stimulus properties such as the force direction (Johansson & Birznieks, 2004). This effect could arise from different patterns and type (contraction vs elongation) of strain wave with respect to motion direction that would preferentially be encoded by tactile afferent. Consistent with previous measurements restricted to the contact, the evolution of the strain wave was stereotypical to the motion direction (Delhaye *et al*., 2016). Furthermore, we show here that the direction of motion created an asymmetrical distribution of the strain with increased total energy on the side of stimulation. However, these patterns were not spatially uniform, and concentration of strains were observed for all participants in particular areas where the skin was compressed.

While surface strains smoothly transitioned across the contact interface and were saliently delineated by the slip region, local curvature changes reflected shifts of the contact area over the fingertip surface. Variations in curvature may therefore provide additional information about contact location and its evolution during manipulation. Such distinct mechanical inputs, surface strain vs curvature, could already constitute partially processed signals that contribute to shaping primary afferent encoding (Platkiewicz *et al*., 2016), consistent with evidence that features of haptic stimuli are represented at early stage of the somatosensory pathway (Bagdasarian *et al*., 2013; Jörntell *et al*., 2014).

### Possible explanation for the wide variability in afferent responses

A recurring observation in neurophysiological studies of tactile is the large apparent variability in afferent response. Wheat *et al*. (2010) reported that “there was no evident relationship between the position of an afferent’s receptive field on the finger” and its firing behavior. In many previous studies, neural responses were pooled across participants using a generic normalized representation of the fingertip. Such approaches may conceal the influence of subject-specific mechanics.

One possible explanation for this variability lies in the complex and highly individualized mechanics of the fingertip. This interpretation is supported by studies showing markedly reduced variability when the mechanical stimulation is precisely controlled in terms of skin strain distributions (Hayward *et al*., 2014) or when strains are measured (Delhaye *et al*., 2021*a*). In the present study, while the global evolution of amplitude of strain energy was largely governed by the slip dynamics, the magnitude and local pattern of deformation were strongly participant-specific, both affected by fingertip size and stiffness.

Deformation patterns were non-uniform over the fingertip surface, particularly outside the contact. Part of this heterogeneity could be explained by the formation of skin wrinkling, i.e. surface irregularities of the skin. Skin wrinkling is a well-known phenomenon in human skin, for example around the eyes and forehead during facial expressions or after prolonged fingertip immersion in water (Yin *et al*., 2010; Wei *et al*., 2025). Our results show that wrinkles can also be formed during tangential loading of the fingertip. Their development is likely influenced by the density and spatial organization of collagen fibers in the dermis (Guissouma *et al*., 2021; Ní Annaidh *et al*., 2012*a*, 2012*b*), which may contribute to individual differences in fingertip mechanical behavior.

## Conclusion

By quantifying strain distributions across the entire fingertip surface, this study reveals that deformation during tangential loading is not confined to the contact interface but extends broadly across surrounding skin regions. The substantial contribution of extra-contact deformation provides a more complete picture of fingertip mechanics during manipulation and highlights the importance of considering whole-fingertip mechanics when interpreting tactile neural responses. These results establish an empirical basis for improving biomechanical models of the fingertip (Duprez *et al*., 2026) and may help bridge the gap between contact mechanics and tactile neural encoding. Future work should further investigate how traction stresses (Li *et al*., 2022) and internal tissue deformation contribute to the mechanical signals driving tactile afferent activity.

## Additional information

### Data availability

All data can be shared upon a reasonable request.

## Acknowledgements

The authors would like to thank Francois Wielant, Julien Lambert and the NTMD team for their technical support. This work was supported by an Incentive Grant for Scientific Research for the Fonds de la Recherche Scientifique – FNRS under Grant(s) No F.4513.25. B.D. is a research associate of the Fonds de la Recherche.

## Supplementary Figures

**Supplementary Figure A1.**
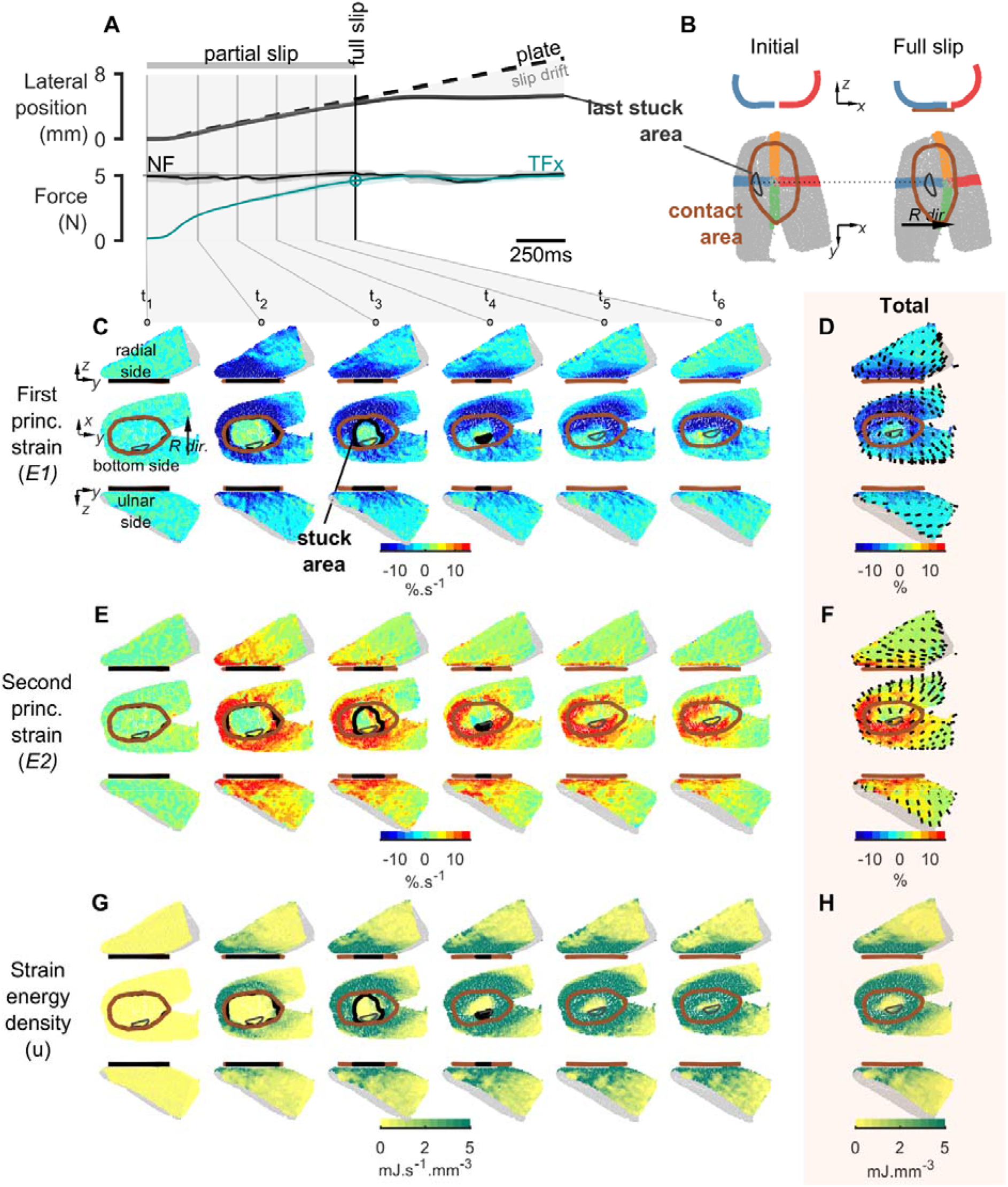
Typical whole fingertip deformation encountered during tangential (radial) loading (S06). (**A**) Evolution of the global variables as a function of time. Top: Lateral position of the plate and of the last part of the skin stuck to the plate. Once the full slip is reached, relative position drift is observed. Bottom: Controlled normal force (black) and tangential force produced in the direction of stimulation (cyan). Solid lines show mean values across repetitions of the same condition, with shaded areas indicating ±1 SD. (**B**) 3D maps of the fingertip reconstruction at initial contact and at full slip. The location of the last stuck region is indicated with crossed bands to help visualize the gross deformation. (**C**) 3D heatmaps of the strain rate evolution during partial slip of the first principal component (E1) at time points defined in (A). Views are shown from the radial side (top), the bottom side (middle), and from ulnar side (bottom). Note that the opposite-facing side is colored in gray. The boundary of the contact area is overlaid in solid brown, the current stuck region in solid black, and the last stuck region is shown in gray. (**D**) 3D heatmaps of total strain (integral of strain rates) for E1, shown in the same perspectives as in (C). Local directions of E1 are overlaid. (**E-F**) Same as (C-D), for the second principal strain component (E2). (**G-H**) Same as in (C-D), for the strain energy density.

**Supplementary Figure A2.**
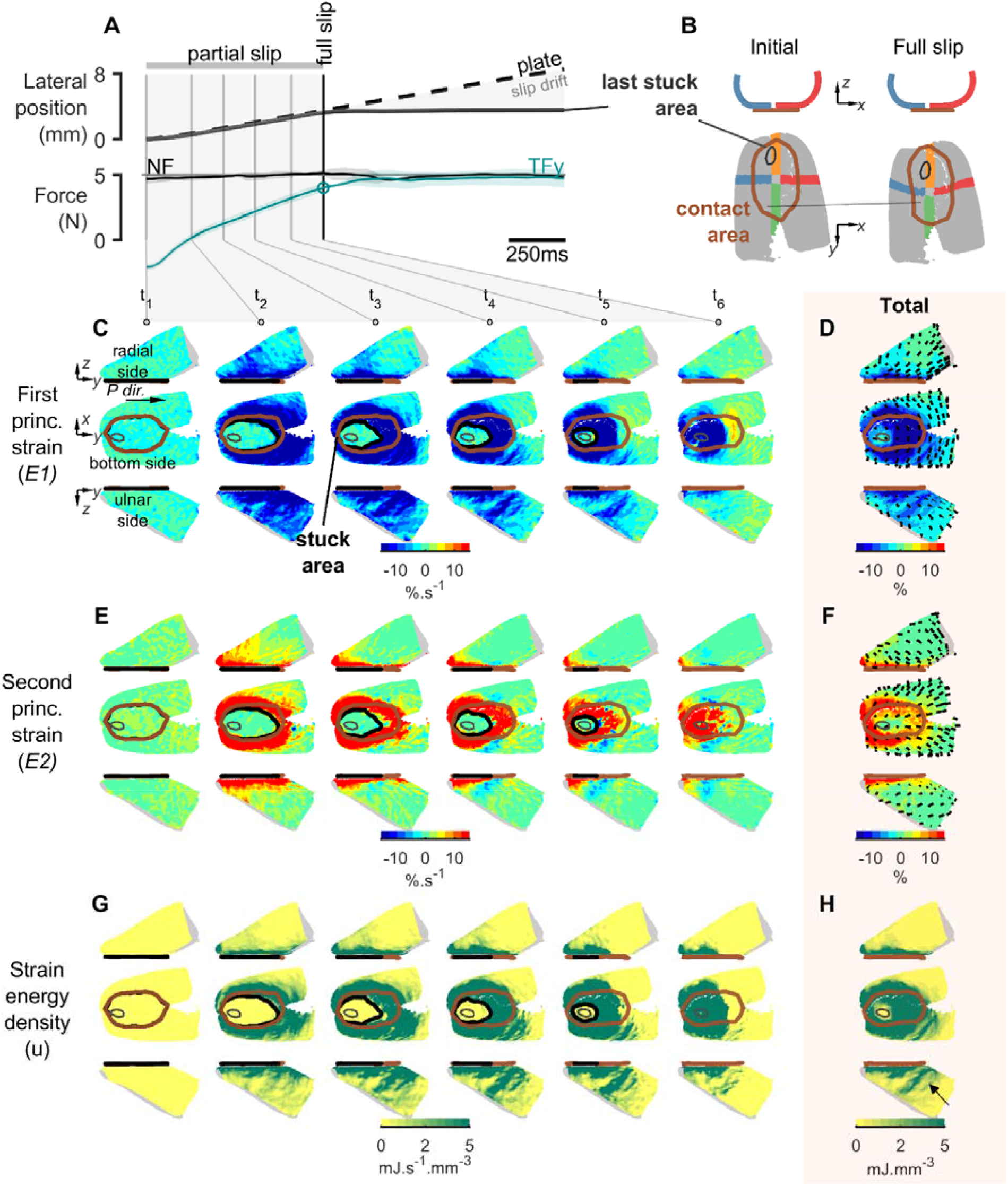
Typical whole fingertip deformation encountered during tangential (proximal) loading (S06). (**A**) Evolution of the global variables as a function of time. Top: Lateral position of the plate and of the last part of the skin stuck to the plate. Once the full slip is reached, relative position drift is observed. Bottom: Controlled normal force (black) and tangential force produced in the direction of stimulation (cyan). Solid lines show mean values across repetitions of the same condition, with shaded areas indicating ±1 SD. (**B**) 3D maps of the fingertip reconstruction at initial contact and at full slip. The location of the last stuck region is indicated with crossed bands to help visualize the gross deformation. (**C**) 3D heatmaps of the strain rate evolution during partial slip of the first principal component (E1) at time points defined in (A). Views are shown from the radial side (top), the bottom side (middle), and from ulnar side (bottom). Note that the opposite-facing side is colored in gray. The boundary of the contact area is overlaid in solid brown, the current stuck region in solid black, and the last stuck region is shown in gray. (**D**) 3D heatmaps of total strain (integral of strain rates) for E1, shown in the same perspectives as in (C). Local directions of E1 are overlaid. (**E-F**) Same as (C-D), for the second principal strain component (E2). (**G-H**) Same as in (C-D), for the strain energy density.

**Supplementary Figure A3.**
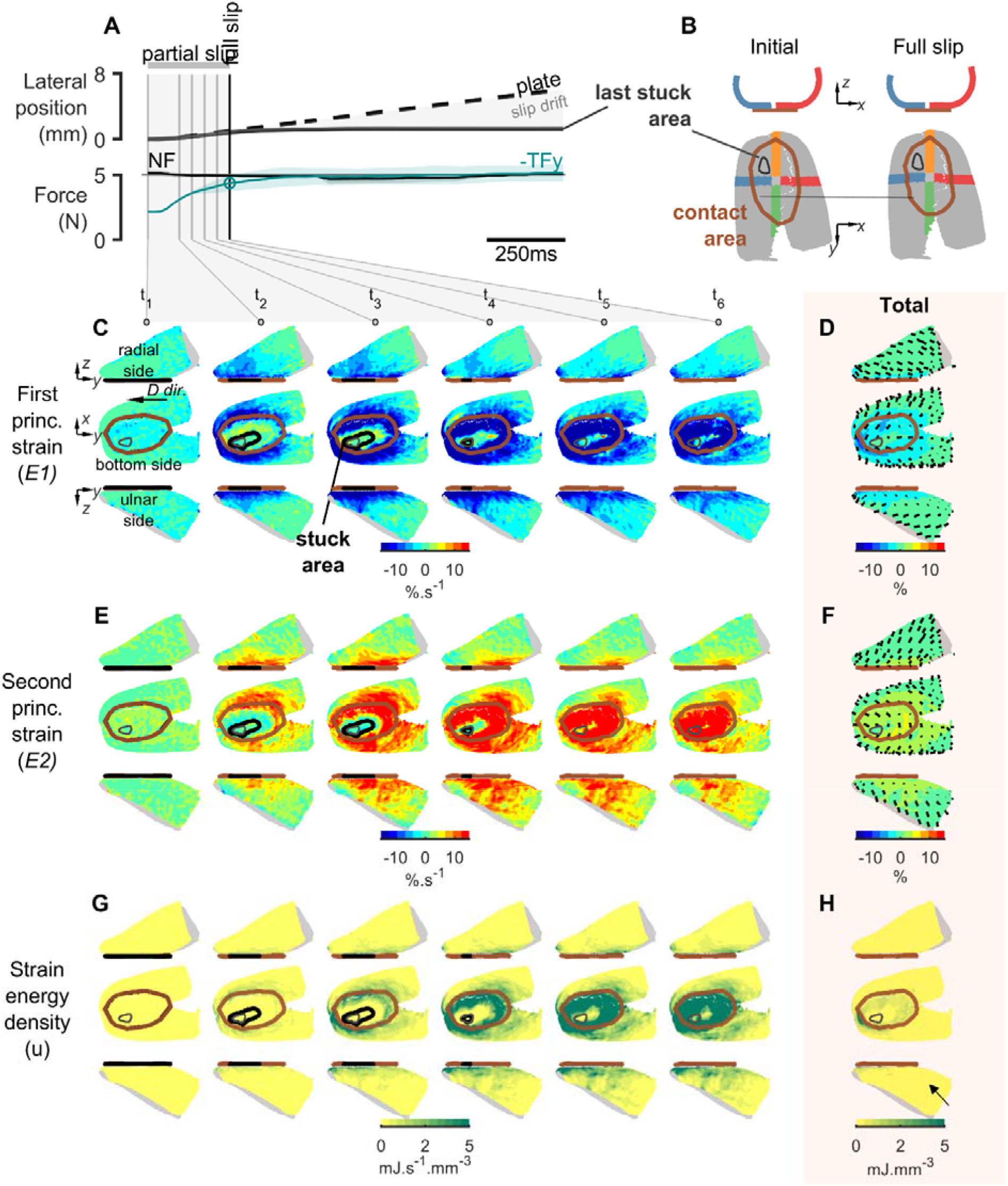
Typical whole fingertip deformation encountered during tangential (distal) loading (S06). (**A**) Evolution of the global variables as a function of time. Top: Lateral position of the plate and of the last part of the skin stuck to the plate. Once the full slip is reached, relative position drift is observed. Bottom: Controlled normal force (black) and tangential force produced in the direction of stimulation (cyan). Solid lines show mean values across repetitions of the same condition, with shaded areas indicating ±1 SD. (**B**) 3D maps of the fingertip reconstruction at initial contact and at full slip. The location of the last stuck region is indicated with crossed bands to help visualize the gross deformation. (**C**) 3D heatmaps of the strain rate evolution during partial slip of the first principal component (E1) at time points defined in (A). Views are shown from the radial side (top), the bottom side (middle), and from ulnar side (bottom). Note that the opposite-facing side is colored in gray. The boundary of the contact area is overlaid in solid brown, the current stuck region in solid black, and the last stuck region is shown in gray. (**D**) 3D heatmaps of total strain (integral of strain rates) for E1, shown in the same perspectives as in (C). Local directions of E1 are overlaid. (**E-F**) Same as (C-D), for the second principal strain component (E2). (**G-H**) Same as in (C-D), for the strain energy density.

